# Ancestral sex-role plasticity facilitates the evolution of same-sex sexual behaviour

**DOI:** 10.1101/2022.06.20.496918

**Authors:** Nobuaki Mizumoto, Thomas Bourguignon, Nathan W. Bailey

## Abstract

Recent attempts to explain the evolutionary prevalence of same-sex sexual behaviour (SSB) have focused on the role of indiscriminate mating. However, in many cases, SSB involves plastically adjusting sex roles to achieve successful courtship or pairing. To evaluate this overlooked factor, we tested whether ancestral sex-role plasticity facilitated the evolution of SSB in the termite *Reticulitermes speratus*. Male termites follow females in paired ‘tandems’ before mating, and movement patterns are sexually dimorphic. Adaptive same-sex tandems occur in both sexes. We show that in such cases, one partner adopts the other sex’s movement patterns, resulting in behavioural dimorphism. Data-based simulations confirmed that this socially-cued plasticity contributes to pair maintenance because dimorphic movements improve reunion success upon accidental separation. Phylogenetic analysis indicated that the ancestors of modern termites lack consistent sex roles during pairing, indicating that *R. speratus* inherited the plasticity from the ancestor. Socio-environmental induction of ancestral behavioural potential may be of widespread importance to the evolutionary maintenance of SSB.

## Introduction

Same-sex sexual behaviour (SSB) is widespread in non-human animals, and its evolution has attracted research interest because of its assumed fitness costs [1–4]. Recent theory has suggested that selection for indiscriminate mating can maintain SSB [5], and ancestral indiscriminate mating has been suggested to underlie the evolutionary origins of SSB [6–8]. However, these arguments overlook an important consideration. SSB is often more than misdirected behaviour otherwise expressed during heterosexual encounters. In many species, sexual interactions between same-sex partners involve at least one partner expressing behaviours associated with the other sex, and evidence is emerging that SSB can involve distinct behavioural repertoires with separate neurological causation [9]. The sex of the partner in a same-sex pairing is different from that in a heterosexual pairing, so behavioural plasticity in response to the immediate sociosexual environment is likely to be integral to the expression of SSB in many cases. Here we use a well-characterised insect system to test whether and how such sex-role plasticity facilitates the evolution of SSB.

In the termite *Reticulitermes speratus*, life-long monogamous pairs establish colonies and produce thousands of offspring [10]. During a brief period, alates (winged adults) disperse from their nests. Both females and males land on the ground, shed their wings, and run to search for a mating partner [11]. Upon joining, a pair performs a tandem run. The male follows the female, maintaining contact in a highly coordinated manner while seeking a suitable site for colony foundation. Tandem running is a form of sexual behaviour as it involves communication via sex pheromones [12], with sex roles being strictly determined: females lead and males follow. A stable tandem run is required for successful reproduction and is maintained via two behavioural processes. First, bidirectional feedbacks between females and males enable them to actively regulate movement speed according to partner distance [13]. Second, sexually dimorphic movements are expressed upon accidental separation, where females pause and males engage in an intensive search to facilitate re-encounter [14]. Same-sex pairing and tandem runs are also observed in *R. speratus*. Termites cannot survive alone, and same-sex pairing enhances both survival and fitness [15,16]. For example, same-sex tandems dilute risk from predators that capture a single prey at a time [17,18]. Female-female pairs can reproduce via parthenogenesis [19], and male-male pairs can invade neighbouring incipient colonies to gain reproductive opportunities [16].

We used *R. speratus* to test the hypothesis that ancestral plasticity in sex roles potentiated the evolution of SSB. Using detailed behavioural analyses of each partner during heterosexual and same-sex pairing, we showed that individuals in same-sex tandems plastically modify their movements to achieve dimorphic behavioural processes similar to that of heterosexual tandems. We then used data-informed simulations to evaluate the fitness consequences of sex-role plasticity during SSB by comparing them with scenarios lacking plasticity. Finally, we determined the origin of sex-role plasticity using a comparative phylogenetic approach. Our findings illustrate how socio-environmental induction of ancestral plasticity can facilitate the evolution of SSB.

## Results

### Phenotypic plasticity of sexually dimorphic behaviour enables same-sex tandem runs

We first evaluated the overall tendency for the formation of different tandem run combinations in *R. speratus*. Alates were collected in March 2021 from Kagoshima, Miyazaki, and Fukui prefectures, Japan. After swarming, we allowed individuals to freely interact in mixed or single-sex groups composed of individuals randomly selected from the same colony. This ensured that any differences in tandem running behaviour reflected plastic responses to the sociosexual environment experienced by each individual in a pair, as opposed to genetic differences. Within these groups, female-male and male-male tandems occurred more commonly than female-female tandems (GLMM, χ^2^_2_ = 16.151, *P* < 0.001, Fig. S1).

Next, we moved a tandem pair from each group to a separate arena to further observe their behaviour. Once a pair formed, same-sex tandems were as stable as heterosexual tandems, with no difference in the time spent in tandem during five-minute observations across pairing combinations (LMM, χ^2^_2_ = 1.868, *P* = 0.393). Same-sex pairs showed behavioural patterns strikingly similar to opposite-sex pairs. For example, leaders and followers in same-sex tandems regulated their motion acceleration in response to changes in interindividual distances (Fig. S2). When interindividual distance increased, the follower sped up while the leader slowed down to avoid separation. When interindividual distance decreased, the follower slowed down while the leader sped up to avoid collision. The leader-follower role was fixed within a pair. In female-male or female-female pairs, we observed no switch of leader-follower roles (0/54 for female-male, 0/45 for female-female pairs), indicating no competition over position. In male-male pairs, however, both males often tried to follow each other forming circles (competition > 5 seconds observed in 11/54 pairs; Supplementary Video 1). However, this rarely resulted in a switch of leader-follower positions (1/54 pairs).

During tandem runs, leaders and followers strongly synchronised their movements with one another, with no significant difference in movement speeds in any pairing combination (LMM, *P* > 0.05, Fig. 1ABDF, Fig. S3A). However, when pairs became accidentally separated, leaders and followers showed distinct movements. In heterosexual tandems that became separated, leader females paused while follower males continued moving, consistent with previous findings [14]. Thus, movement speed following separation differed notably between leaders and followers in heterosexual pairs (LMM, χ^2^_1_ = 41.246, *P* < 0.001, Fig 1AC, Fig. S3B). We found similarly striking dimorphic movement patterns in separated same-sex tandems: movement patterns were dictated by leader-follower position, not sex. For example, in female-female tandems, leader females slowed down in the manner of females of heterosexual tandems, but follower females kept moving to search for the stray partner in the manner of males in heterosexual tandems (LMM, χ^2^_1_ = 60.471, *P* < 0.001, Fig 1E, Fig. S3B). Similarly, in separated male-male tandems, follower males moved to search for the stray partner while leader males paused (LMM, χ^2^_1_ = 71.077, *P* < 0.001, Fig 1G, Fig. S3B).

**Figure 1.**
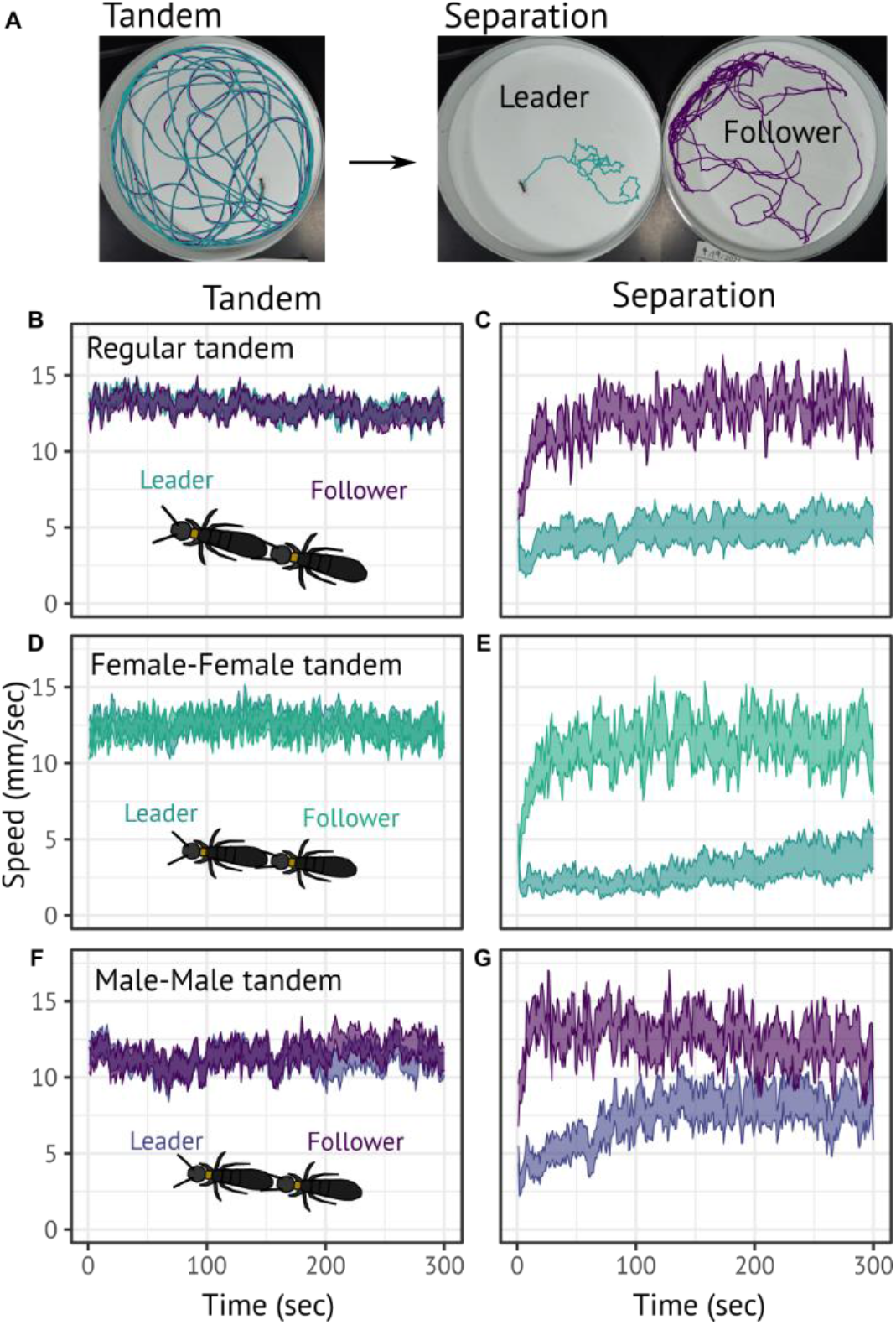
Movement speed of termites in heterosexual or same-sex tandem pairings. (A) Representative five-minute trajectories during tandem run and after separation in heterosexual pairs. (B-G) Mean movement speed of individuals during tandems (B, D, F) and after separation (C, E, G). One individual was removed from each tandem at time = 0 in the right column to observe behaviour after separation. Shaded regions indicate mean ± 1 S.E.

Male and female same-sex pairings did not generate identical behavioural dynamics, however. After separation, females in female-female pairs moved slower than males in male-male pairs (LMM, Tukey’s test, *P* < 0.05 between female-female pairs and male-male pairs both in leaders and followers, Fig. S3B). Also, leaders and followers showed distinct turning patterns after separation irrespective of pairing combinations, with leaders showing higher movement sinuousity than followers after separation (LMM, *P* < 0.001). These results indicate that females and males in same-sex tandems plastically adjust to their sociosexual environment, modifying their behaviours to achieve movement coordination as effective as that of heterosexual tandems.

### Dimorphic movement enhances the likelihood of tandem reunion in separated same-sex pairs

In heterosexual tandem runs, sexually dimorphic movements after separation are key to efficient re-encounters [14]. We tested whether the dimorphic movements observed in same-sex pairs enhance the probability of reunion after separation using simulations of an individual-based model. In random search, different movements result in different encounter rates [20]. If male-behaving females or female-behaving males increase the likelihood of reunion in same-sex pairs, then dimorphic movements observed in same-sex pairs are predicted to increase search efficiency compared to scenarios where both females paused or both males continued moving. Using parameters for speed and turning angles estimated from the analyses above, we modelled movement patterns using a correlated random walk (CRW). Then, we evaluated five combinations of movement patterns: the observed dimorphic movements in heterosexual pairs, female-female pairs, and male-male pairs, and ‘virtual’ monomorphic movement scenarios where both partners moved like female leaders or male followers.

Behavioural dimorphism in same-sex tandems increased reunion probability upon separation. In simulated male-male and female-female pairs showing dimorphic movements, the re-encounter probability was comparable to that of heterosexual pairs (Figs 2A-B). In contrast, if both males of same-sex pairs kept their behaviour unchanged (*i*.*e*. both males moved to search for the stray partner), re-encounter probability decreased (Fig. 2A). The same was true for female-female pairs (Fig. 2B), for which re-encounter probability strongly decreased when both females kept their behaviour unchanged (*i*.*e*. both females paused to wait for the stray partner). Thus, the behavioural changes observed in same-sex pairings of termites are critically important to maintain pair coordination.

**Figure 2.**
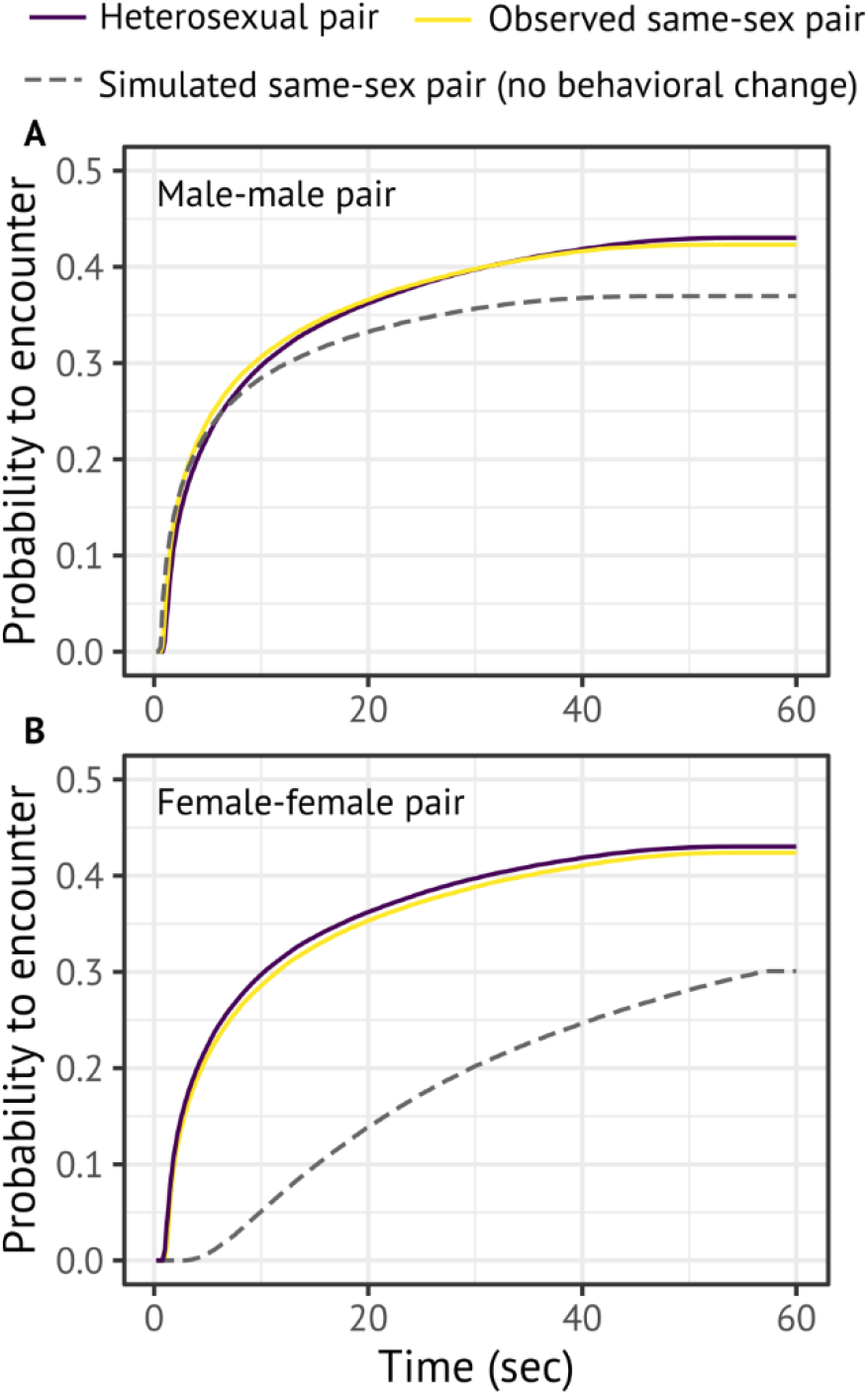
Simulation results for re-encounter efficiency of termite movements after pair separation. Encounter efficiency was compared across CRWs parameterised with observed movements in heterosexual pairing, observed movements in same-sex pairing, and simulated movements involving no behavioural change in same-sex pairing. (A) In cases where one male in a male-male tandem expresses female-like movements upon separation, the resulting movement dimorphism enhances re-encounter rates. (B) Similarly, when one female in a female-female tandem expresses male-like movement, the resulting movement dimorphism enhances encounter rates. Movements were modelled using CRWs with parameters in Table S1, and results were obtained from 100,000 simulations.

### Plasticity of dimorphic behaviours required for adaptive same-sex tandems is ancestral

The above results demonstrate that both female and male termites possess a full behavioural repertoire for mate pairing, and can functionally express behaviours associated with the other sex. We hypothesised that the sex role flexibility observed in *R. speratus* was inherited from the common ancestor of modern termites. Previous studies have implied that sex roles are flexible in early divergent lineages [21], while in Neoisoptera, a clade containing *R. speratus*, females develop tergal glands producing sex pheromone and predominantly perform leader roles [22,23]. We tested this hypothesis by performing an ancestral state reconstruction on the termite phylogenetic tree.

The focal species of this study, *R. speratus*, shows female-leader tandem runs in heterosexual pairs. All other taxa for which we found records in the Neoisoptera also show female-leader tandem runs (n = 21 genera). Our ancestral state reconstruction indicated a 99.97% probability that tandem runs were present in the ancestor of Neoisoptera, and a 99.4% probability that those tandem runs were female-led. A separate reconstruction for males found a negligible probability (2.73%) of male-led tandems in the ancestor of Neoisoptera (Fig. 3).

**Figure 3.**
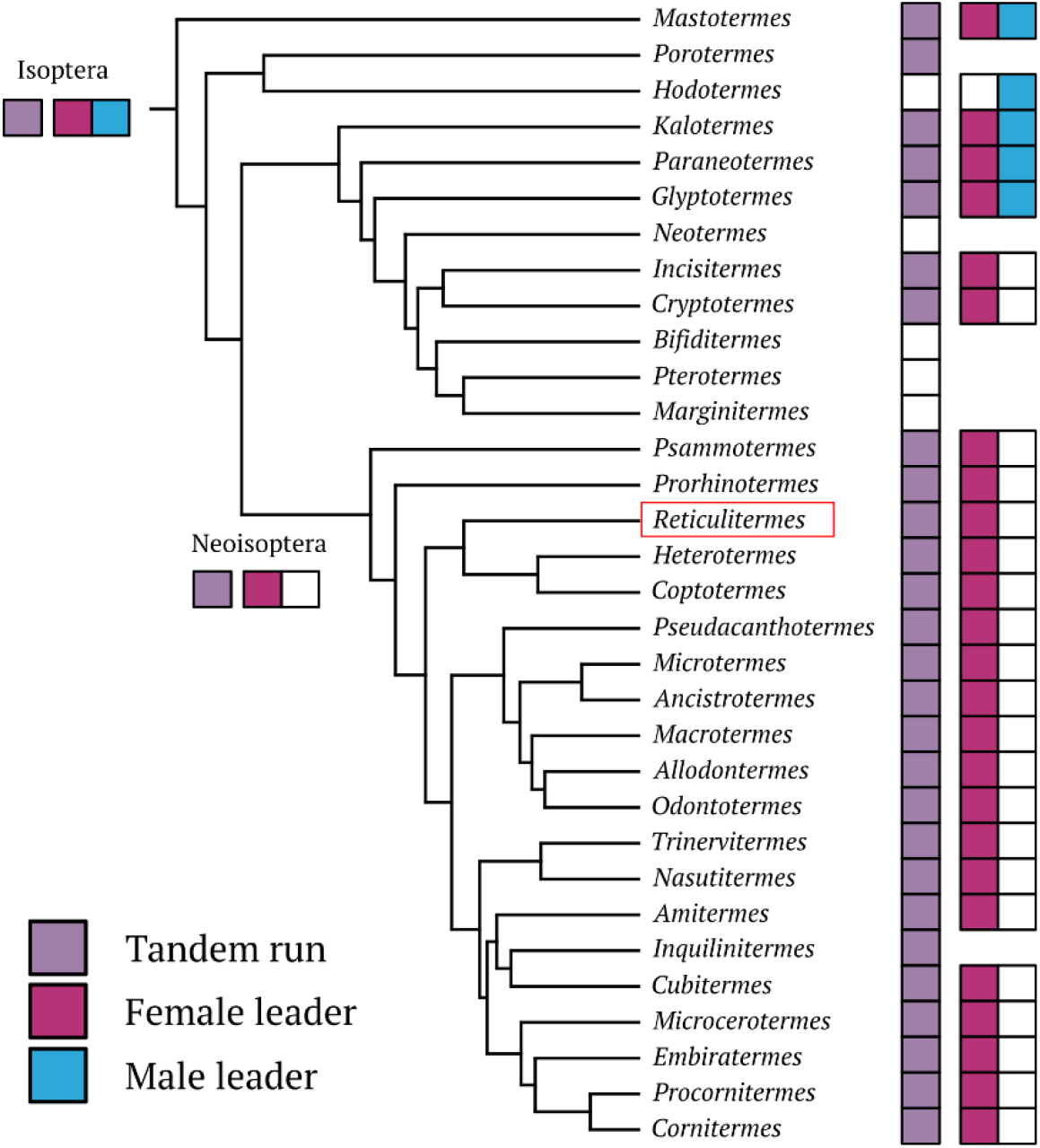
Phylogeny of termites showing tandem run and leader role character states. Inferred ancestral states are shown in the ancestor of Isoptera and Neoisoptera. The genus *Reticulitermes*, used in behavioural observations during this study, is framed by a red rectangle. The absence of squares next to taxa indicates an unknown state.

In notable contrast, sex roles are more flexible in basal termite lineages, and tandem runs may be led by either females or males (Fig. 3). We found that the presence of tandem runs is likely ancestral in termites (98.81% probability). However, there was a greater than 90% probability that both females and males expressed leader roles in the ancestor of modern termites (99.16% for females, and 91.17% for males; Fig. 3). Thus, before females acquired a fixed leader role in the heterosexual tandem runs of Neoisopteran termites, the leader-follower role was flexible. Both females and males were capable of expressing the full behavioural repertoire for movement coordination. These results support the hypothesis that behavioural plasticity was ancestral in termites and was co-opted to facilitate adaptive same-sex tandem runs in Neoisoptera, to which *R. speratus* belongs.

## Discussion

Efforts to identify factors promoting the evolution of same-sex sexual behaviour have recently focused on the role of incomplete sex discrimination [5,8], particularly in arthropod taxa [12,24]. Our results highlight a potentially important alternative that has received less attention: sex role plasticity. In the termite *R. speratus*, plasticity of sexually dimorphic behaviour during tandem running is necessary for successful same-sex pair coordination in both sexes. In heterosexual tandems, the leader-follower role is usually associated with sex. Leader females pause to wait if they are separated from males, while follower males engage in an intensive search for their partner upon separation (Fig. 1A-C). However, we found that both females and males retain the behavioural potential to flexibly adopt the role of the other sex when they interact in same-sex tandems (Fig. 1D-G). Simulations parameterised with experimental data showed that sensitivity to sociosexual context and role flexibility are important for maintaining same-sex tandems (Fig. 2), and comparative phylogenetic analysis indicated sex role plasticity is ancestral in termites. Consistent with this, some non-Neoisopetra termites, such as *Mastotermes, Paraneotermes*, and *Glyptotermes*, have no sex-associated leader-follower roles, and both sexes are capable of expressing either behaviour (Fig. 3). Our results suggest that ancestral behavioural plasticity played an essential role in the evolution of same-sex sexual behaviour. Also, the results provide new insight into the relationship between SSB and other sexual traits by highlighting important considerations about the differences between female and male SSB.

A comparison between termite same-sex pairing and SSB associated with other mating systems provides useful insight into the evolution of mating behaviour diversity. In same-sex termite pairs, at least one individual expresses behaviours associated with the other sex in the context of heterosexual pairing. Such behavioural mimicry of the other sex is known to evolve as an alternative mating tactic in many taxa. For example, when the density of competitors is very high in many insects, birds, and reptiles, males mimic females to avoid intraspecific competition [25,26] or prevent others from copulating with females [27–29]. Similarly, females may mimic males to reduce the cost of copulation or male harassment [30]. Such sexual mimicry might coevolve with SSB because SSB can also reduce aggressive interactions during competition [31,32].

SSB might differ both qualitatively and quantitatively between females and males, as many species develop sexually differentiated behavioural, signalling, and life-history traits [33,34]. Our study uncovered a sex difference in same-sex pair coordination in *R. speratus*. First, males were more likely to form same-sex tandems (Fig. S1) and compete to obtain follower positions, whereas females in same-sex tandems showed no such competition. To initiate tandem runs, at least one individual needs to begin following another individual. The implications of this are significant. As females are the leaders in heterosexual tandems, one female must switch her behavioural strategy before forming a same-sex tandem. The factors that cue this anticipatory response are uncertain, but may involve integrating information about the encounter rate of either sex over successive interactions. In contrast, males in heterosexual tandems always adopt follower roles, meaning males can initiate same-sex tandems before they establish who takes the leader position. This asymmetry explains why females took longer to initiate same-sex tandems than males. Second, after separation, leader males quickly stopped waiting for the stray same-sex partner and moved away to search for another potential mate, while leader females showed lengthy and stable pausing behaviour (Fig. 2). This difference could be explained by the pair-bonding pheromones females secrete to attract males from a short distance [22]. In random search with sexual attraction signals, the signalling sex achieves higher encounter rates by moving slower or even pausing, while the sex without signals should move actively to enhance encounters [35,36]. Thus, pausing is not suitable for male termites if they cannot re-encounter the stray partner in a short period, while for females, pausing can be beneficial for re-encounters or encounters with new potential mates due to the effects of their mate-attraction signals.

The loss of sexual signals is often thought necessary for the evolution of same-sex behaviour [5,37], because without strong sexual signalling, or the ability to perceive and differentiate sexual signals, individuals may not distinguish the sex of the mating partner and thereby engage in same-sex behaviour [38]. However, our results emphasise that same-sex pairing can be maintained even with strong sexual signals. Neoisopteran termites, including *R. speratus*, express sex-associated leader-follower roles in heterosexual tandems, with females the leaders and males their followers (Fig. 3). In this clade, only females have sex-pairing pheromones, while in basal lineages, both females and males emit pairing pheromones [22,39]. Thus, the evolution of sexually dimorphic behaviour in tandems is clearly associated with the evolution of sex-specific signals, strongly arguing against a “mistaken identity” hypothesis of same-sex pairing in termites.

Recent theoretical work has proposed that SSB can be evolutionarily maintained through selection for indiscriminate sexual behaviour [5], and a conceptual model has argued that the prevalence of SSB across widespread animal taxa might be due to inheritance of incomplete sex discrimination from a common ancestor of sexually reproducing species [8]. However, these explanations for the prevalence of SSB in animals are based only on the viewpoint of active mates (*i*.*e*. interacting partners that attempt to court, mount, pair, or copulate with one another). The perspective of passive mates (*i*.*e*. interacting partners being courted, mounted, paired, or receiving a copulation attempt) is often lacking, and as a result, it is underappreciated that both individuals may modify their behaviour during SSB to maintain same-sex pairing [40]. In this study, we empirically demonstrated that one of the partners in a same-sex pairing of termites flexibly expresses the behaviour of the other sex to contribute to pair coordination. Importantly, accurate sex discrimination, rather than failures of it, and induction of plastic responses to the sociosexual environment are both prerequisites for the stable occurrence of SSB. Therefore, it is important to expand our focus from incomplete sex discrimination to the diversity of factors that facilitate the evolutionary origins of SSB, particularly in taxa, where sexually dimorphic interactions are involved.

## Materials and Methods

### Behavioural observations

We collected *R. speratus* alates with a piece of nesting wood from five colonies in Kagoshima (colonies A-B), Miyazaki, and Fukui (colonies E-F) in March 2021, just before their swarming season. To control flight timing, all nesting wood was maintained at 22°C until experiments began. Before each experiment, we transferred nests to a room at 27°C, which promoted alates to emerge and fly. Alates were then collected and separated individually. We used individuals that shed their wings by themselves within 12 hours.

The frequency of tandem runs was likely to vary among different pair combinations. To ensure that we observed pair coordination for each combination equally, we first prepared a source group for tandem runs and then extracted a pair for further observations. Three source groups were used to generate heterosexual, female-female, and male-male tandems, respectively consisting of 1) five females and five males, 2) ten females, or 3) ten males. In each case, the 10 individuals were placed in a petri dish (ø=140mm) with moistened plaster. All individuals were marked with one coloured dot of paint (PX-20; Mitsubishi) on the abdomen to distinguish individual identity. Groups were maintained for more than 30 minutes to ensure tandem formation. Each group was recorded with a video camera (HC-X1500-K, Panasonic), and we counted the number of tandem running individuals at 15 minutes. The number of tandem running individuals was compared across treatments using a generalised linear mixed model (GLMM) with binomial distribution and logit link, where “original colony” was treated as a random effect. All members within the same source groups were from the same original colony.

We transferred a single tandem pair to an observation arena consisting of a petri dish (ø=90mm) with moistened plaster. The tandem was disturbed by transportation, but most pairs restarted tandems after being introduced to the new arena. Once the tandem run resumed, we recorded behaviour of the pair for five minutes using a video camera as above. After five minutes, we carefully removed one individual using an aspirator and observed the movement of the separated individual during reunion search (“separation search”). Each individual was used only once for data collection. We obtained 55 observations for heterosexual tandems (6, 13, 14, 8, and 14 for colonies A, B, C, E, and F, respectively), which included observations of separation search by 29 leader-females and 26 follower-males. Similarly, we observed 46 female-female same-sex tandems, with 22 leader-female and 24 follower-female observations of separation search, and 56 male-male same-sex tandems, with 29 leader-male and 27 follower-male observations of separation search. We extracted the coordinates of termite movements from all videos using the video-tracking system UMATracker [41]. We allocated “id 0” to the individual that was the leader at the beginning, and “id 1” for the follower, and manually checked all videos to verify that leader-follower roles were not inadvertently flipped during coordinate extraction. We down-sampled all videos to a rate of five frames per second (FPS) (= every 0.2002 s) for subsequent analyses. All data analyses were performed using R v. 4.0.1 [42]. The source codes for data analysis are available in Data S2.

### Tandem analysis

Our first analysis tested whether behaviour differs in same-sex versus heterosexual tandems. To compare the time engaged in tandem runs across pairs, we automatically identified whether a pair was performing tandem runs for every video frame, combining methods described in previous studies [14,43,44]. During observations, pairs were determined to be in one of three states: (i) tandem running, (ii) interacting but not tandem running, and (iii) searching (individuals in the pair are physically separated). We defined individuals in the pair as interacting (or tandem running) when the distance between their centroids was less than 7mm [14]. This distance slightly exceeds the average body length because termites in a tandem run are nearly in physical contact [14]. An interacting pair was considered to be performing a tandem run only if they met the following criteria [43]. First, the interaction needed to last for more than 2 seconds; a very short separation (< 2 seconds) was not regarded as a separation event. Second, both termites needed to move more than 30 mm while interacting. After separation, we considered that individuals were engaging in separation search until they interacted again for more than 1 second.

In male-male tandem runs, the two males occasionally chased each other, formed a small rotation, and competed over the follower position (Supplementary Video 1). We considered this competition state as a special case of tandem runs. The competition state was automatically defined as follows. First, the distance between individuals needed to be smaller than 4mm as two individuals in such a state were located side by side for their heading direction and facing the opposite direction (Supplementary Video 1). Second, the rotation index of a pair needed to be larger than 0.5. The rotation index is calculated as the sum of the angular momenta about the centre of the pair, taking values between 0 (no rotation) and 1 (strong rotation) [45]. Finally, as in tandem runs, the above two conditions needed to persist for more than 2 seconds. These thresholds were determined arbitrarily according to our visual inspection, but modification did not change our conclusions.

We compared the proportion of time performing tandem runs across different pair combinations using a linear mixed model (LMM), with pair combination (heterosexual, female-female, or male-male) treated as a fixed effect and the original colony as a random effect. We transformed proportional data using logit-transformation after adding 0.01 to the observed proportions to avoid infinite values [46]. A likelihood-ratio test was used to test for statistical significance of the inclusion of each explanatory variable (type-II test).

### Movement analysis upon separation

In heterosexual tandems, a pair shows sexually dimorphic movements after separation in which leader females pause while follower males move to enhance reunion [14,43]. To investigate how termites in same-sex tandems behave upon accidental separation events, we computed displacements of individual positions for every frame during tandem runs and after artificial separation. Then, we summed up five successive displacements to obtain moved distance per second as a proxy for movement speed (mm/sec). We focused on the last 1min for tandem runs and the first 1min after separation to obtain mean movement speed for each individual so that we could compare values across all combinations of sexes, roles, and contexts. These mean speeds were compared between leaders and followers for each pairing combination, using an LMM treating leader-follower role as a fixed effect and the original colony as a random effect. Also, we compared speed among pairing combinations for leader and follower separately, with an LMM treating pairing combinations as a fixed effect and the original colony as a random effect. In models where the pairing combination was significant, we ran a Tukey’s post hoc test using the glht() function in the package “multcomp”.

In addition to movement speed, we analysed sinuosity (turning patterns) of termite movements within the same time windows. We computed turning angle as the magnitude of change in the direction of motion from one frame to the next frame. Then, we fit wrapped Cauchy distributions to turning angle data for each individual, using maximum likelihood estimation methods, and took the distribution’s scale parameter as a proxy of sinuosity [47]. Depending on the value of the scale parameter, the wrapped Cauchy distribution varies from a uniform distribution (scale parameter = 0, maximum sinuosity Brownian walk) to a delta distribution (scale parameter = 1, minimum sinuosity straight walk). We compared the value of the scale parameter using an LMM as in the movement speed analysis. Although these parameters did not always follow normal distributions, LMM is robust against violations of distribution assumptions [48].

### Individual-based model

We developed an individual-based model to test whether the observed behavioural dimorphism in separation search movement enhances reunion efficiency in same-sex pairs of termites, to the same extent that it does in heterosexual pairs. When termites in a pair are accidentally separated, the two individuals should be close to each other, but they are not certain where the separated partner is. This situation can be simulated by expressing where two random searchers are located when at distance *d* in a borderless two-dimensional continuous space [14]. The distance *d* was obtained from observed separated distances during spontaneous separation in the tandem observations above. Our simulation considered random searchers walking until re-encountering a partner. An encounter was regarded to have occurred when the distance between the centres of the two individuals became smaller than φ. The value φ was based on our definition of tandem running (=7 mm).

Individuals performed a correlated random walk (CRW) with parameters of speed and sinuosity, denoted *v* and *ρ*, respectively. The speed parameter *v* was obtained as the mean value of the movement speed for each sex and role during the 1 min after separation, while the sinuosity parameter *ρ* was obtained as the estimated scale parameter from the data of turning angles. Parameter values are summarised in Table S1. Based on our behavioural analysis, each time step was adjusted to 0.2s. Thus, each individual moved 0.2*v* mm in each time step. Turning angles followed wrapped Cauchy distribution with scale parameter *ρ*. After generating a uniform random number *u* (0 < *u* ≤ 1), the turning angles θ were derived from the following equation by applying the inversion method [47]:

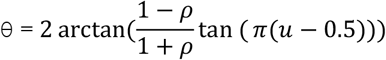

We initiated the simulation with a random bearing angle that fluctuated according to θ. At each step, the bearing angle was equal to the previous bearing angle plus the deviation θ such that the moving object always kept approximately the previous direction, forming a CRW. Movement models can be more complex, adding move/pause patterns, temporal changes of moving speed, initial heading directions, etc. However, previous studies show that movement speed is the most important parameter, and these additional complexities do not change general conclusions about termite reunion efficiency [14,43].

We compared the searching efficiency among five possible combinations of movement patterns. Simulations were performed for 60sec (= 300 time steps). We ran 100,000 simulations and measured efficiency as the probability the focal individual encounters their partner. Simulations were implemented in Microsoft Visual Studio C++ 2019. The source codes for all simulations are available in the Data S3.

### Phylogenetic comparative analysis

We assembled information about termite tandem runs across 137 extant genera present in the phylogeny used in [49]. This phylogeny was inferred from previously published complete mitochondrial genome data combined with morphological characters. Note that this phylogeny randomly selected one representative species for every genus, and thus the polyphyletic genera, such as *Nasutitermes*, were represented as monophyletic groups occupying singular phylogenetic positions. We pruned extinct species from the phylogenetic tree because no information about tandem runs is available.

We used Google Scholar to collect literature data on termite tandem runs, by searching for every genus name with the query “tandem.” In every genus, we compiled the record of tandem run observations with the information about which sex(es) play the leader role. In *Mastotermes darwiniensis*, although there was no information about tandem runs by mating partners, workers perform tandem runs to recruit nestmates to new food resources [50]. We classified this species as performing tandem runs, and both females and males can take the leader role as workers are sex-less individuals. *Hodotermes mossambicus* does not show explicit tandem runs, but males emit pheromones to attract females to follow them [21]. Thus, we treated this species as non-tandem running, but with males expressing a leader role. In addition to the literature data, we counted a termite genus, *Heterotermes*, as performing female-leader tandem runs based on personal observations (N. Mizumoto) of *Het. aureus* after monsoon rainfall in Tempe, Arizona, 2019. We also counted *Glyptotermes* as performing tandem runs during which both females and males perform a leader role based on personal observations (N. Mizumoto) of *Gly. fuscus* (collected in Okinawa) and *Gly. satsumensis* (collected in Kagoshima) under laboratory conditions. We verified that excluding the genera with personal observations did not change our conclusion by re-running comparative analyses without them. The results were qualitatively similar and quantitatively nearly identical except for the ancestral state of male leader in Isoptera (68.6%). In total, we obtained data from 32 genera with phylogenetic information (Data S1).

Then, we carried out separate ancestral state reconstructions for tandem running, female leadership, and male leadership, respectively, using the function ace() in the R package “phytools” [51]. We used a maximum likelihood model with an equal rate of transition among states.

## Acknowledgements

We thank Ales Bucek, Kensei Kikuchi, and members of Bailey lab for helpful discussion; Kensei Kikuchi and Simon Hellemans for termite collection. This study is supported by a JSPS Research Fellowships for Young Scientists CPD (grant number: 20J00660) and a Grant-in-Aid for Early-Career Scientists (21K15168) to N.M., and OIST core funding. N.W.B. gratefully acknowledges funding from the UK Natural Environment Research Council (NE/T000619/1).

## Authors contributions

N.M.: conceptualisation, data curation, formal analysis, funding acquisition, investigation, methodology, resources, validation, visualisation, writing-original draft, writing-review & editing. T.B.: resources, writing-review & editing. N.W.B.: conceptualisation, writing-original draft, writing-review & editing.

## Data accessibility statement

All data will be available in Figshare.

---

**Figure S1.**
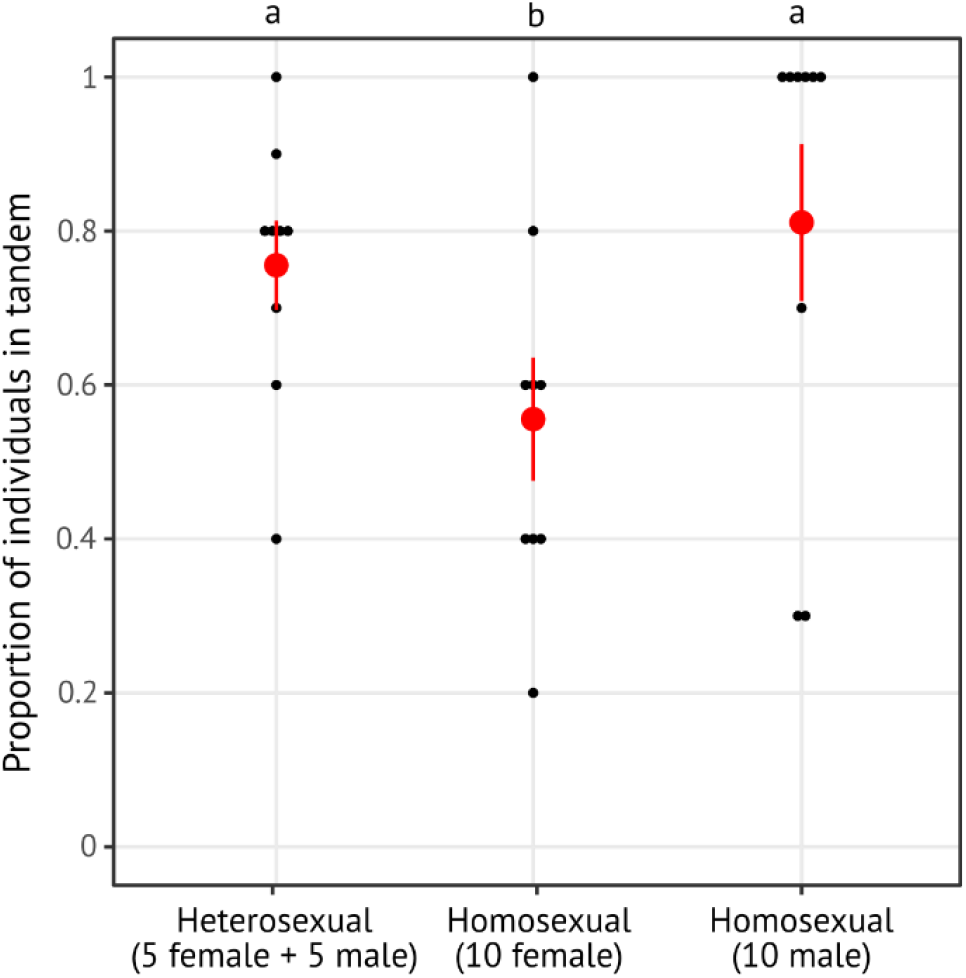
Comparison of the tendency to form tandems among heterosexual and same-sex combinations. Red points and bars indicate means ± 1 s.e. Different letters indicate a significant difference (GLMM with Tukey’s test, *P* < 0.05).

**Figure S2.**
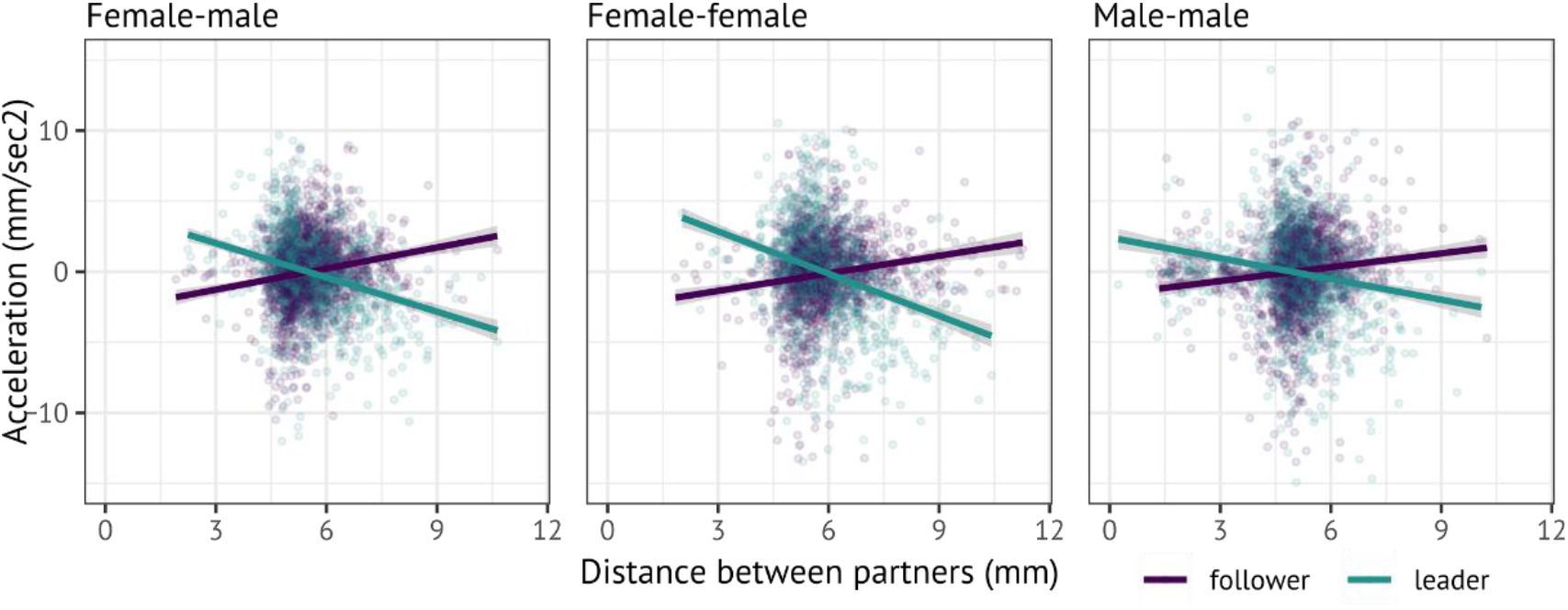
Regulation of motion accerelation during tandem runs. Plots were reduced to 10,000 data points randomly selected from each whole dataset. Slopes were significantly greater than 0 in followers (LMM with “colony” as a random effect, *P* < 0.001, female-male: estimate = 0.535, female-female: estimate = 0.473, male-male: estimate = 0.300), while slopes were smaller than 0 in leaders (LMM with “colony” as a random effect, *P* < 0.001, female-male: estimate = -0.845, female-female: estimate = -1.22, male-male: estimate = -0.480).

**Figure S3.**
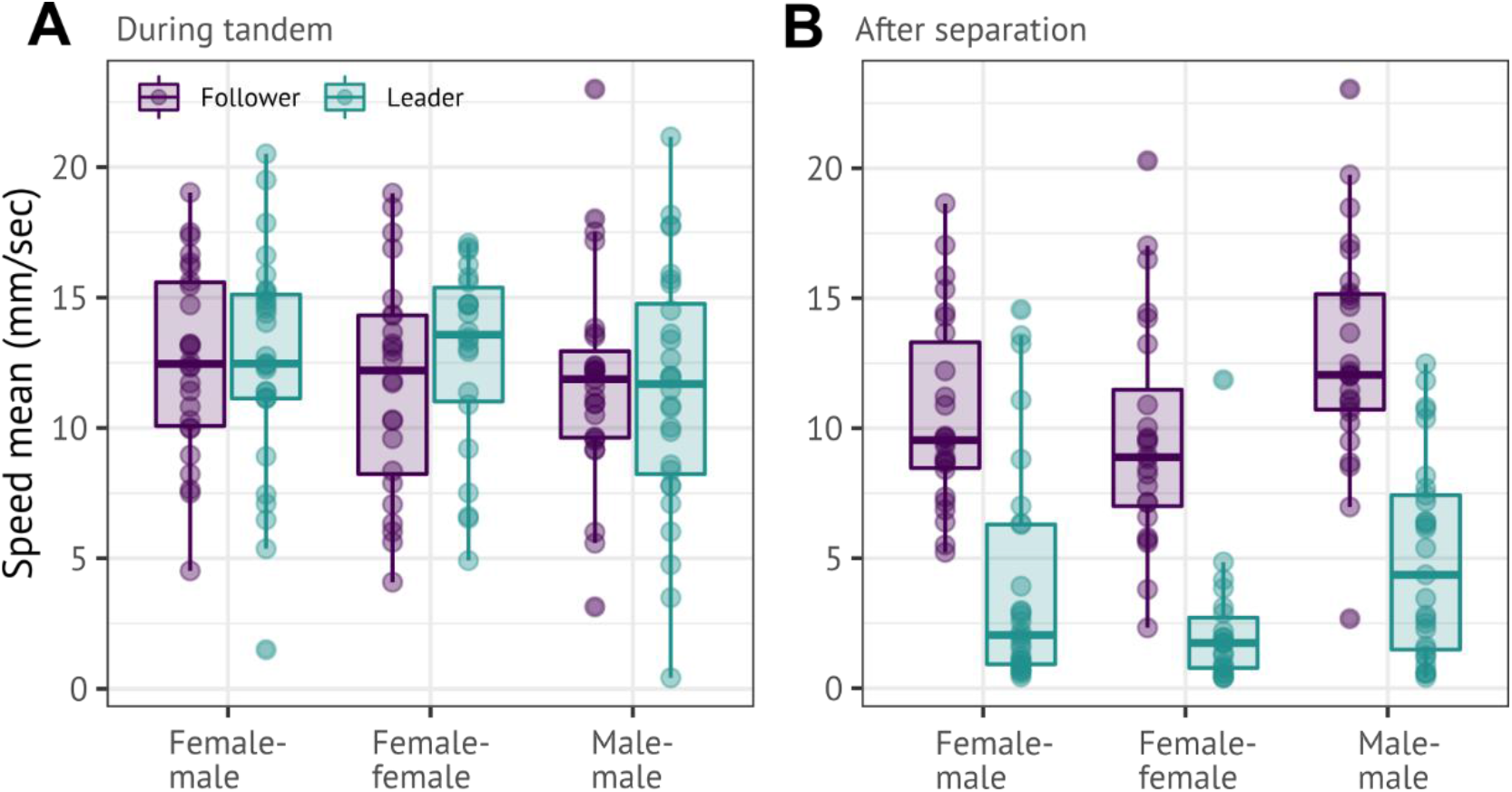
Comparison of mean movement speeds. Speeds during tandem runs (A) show data from the last 1 min of a 5 min observation period, and speeds after separation (B) show data from the first 1 min after separation. Boxplots show quartiles and range (excluding points that are below the lower quartile by at least 1.5 times the interquartile range). All data points were plotted as dots.

**Table S1.**
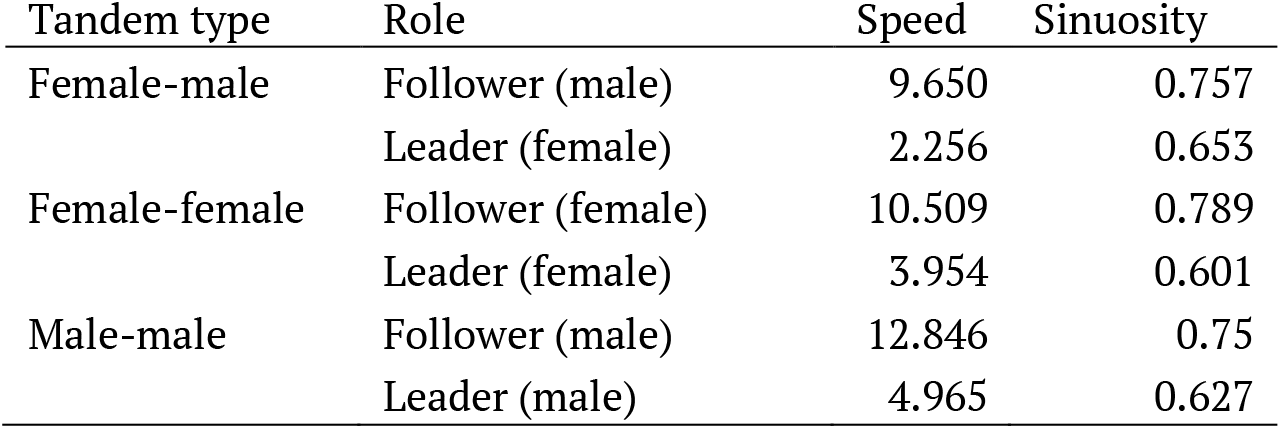
Parameters for correlated random walk used in simulations.

| Tandem type | Role | Speed | Sinuosity |
| --- | --- | --- | --- |
| Female-male | Follower (male) | 9.650 | 0.757 |
|  | Leader (female) | 2.256 | 0.653 |
| Female-female | Follower (female) | 10.509 | 0.789 |
|  | Leader (female) | 3.954 | 0.601 |
| Male-male | Follower (male) | 12.846 | 0.75 |
|  | Leader (male) | 4.965 | 0.627 |

**Supplementary Video 1.** Representative competition rotation by male same-sex tandem pair in *Reticulitermes speratus*.

## Notes

### Competing Interest Statement

The authors have declared no competing interest.

## References

1. Bagemihl B. 1999 Biological exuberance: Animal homosexuality and natural diversity. New York: NY: St. Martins’ Press. See https://scholar.google.co.jp/scholar?q=Biological+Exuberance:+Animal+Homosexuality+and+Natural+Diversity+&hl=ja&as_sdt=0,5#0.

2. Sommer V, Vasey PL. 2006 Homosexual behaviour in animals: an evolutionary perspective. Cambridge: UK: Cambridge University Press.

3. Bailey NW, Zuk M. 2009 Same-sex sexual behavior and evolution. Trends Ecol. Evol. 24, 439–446. (doi:10.1016/j.tree.2009.03.014)

4. Poiani A. 2010 Animal homosexuality: a biosocial perspective. Cambridge: Cambridge University Press.

5. Lerch BA, Servedio MR. 2021 Same-sex sexual behaviour and selection for indiscriminate mating. Nat. Ecol. Evol. 5, 135–141. (doi:10.1038/s41559-020-01331-w)

6. Clive J, Flintham E, Savolainen V. 2020 Understanding same-sex sexual behaviour requires thorough testing rather than reinvention of theory. Nat. Ecol. Evol. 4, 784–785. (doi:10.1038/s41559-020-1189-3)

7. Dickins TE, Rahman Q. 2020 Ancestral primacy of same-sex sexual behaviour does not explain its stable prevalence in modern populations. Nat. Ecol. Evol. 4, 782–783. (doi:10.1038/s41559-020-1187-5)

8. Monk JD, Giglio E, Kamath A, Lambert MR, McDonough CE. 2019 An alternative hypothesis for the evolution of same-sex sexual behaviour in animals. Nat. Ecol. Evol. 3, 1622–1631. (doi:10.1038/s41559-019-1019-7)

9. Karigo T, Kennedy A, Yang B, Liu M, Tai D, Wahle IA, Anderson DJ. 2020 Distinct hypothalamic control of same-and opposite-sex mounting behaviour in mice. Nature 589. (doi:10.1038/s41586-020-2995-0)

10. Vargo EL, Husseneder C. 2009 Biology of subterranean termites: insights from molecular studies of Reticulitermes and Coptotermes. Annu. Rev. Entomol. 54, 379–403. (doi:10.1146/annurev.ento.54.110807.090443)

11. Nutting WL. 1969 Flight and colony foundation. In Biology of termites (eds K Krishna, FM Weesner), pp. 233–282. New York: Academic Press. (doi:10.1016/B978-0-12-395529-6.50012-X)

12. Scharf I, Martin OY. 2013 Same-sex sexual behavior in insects and arachnids: prevalence, causes, and consequences. Behav. Ecol. Sociobiol. 67, 1719–1730. (doi:10.1007/s00265-013-1610-x)

13. Valentini G, Mizumoto N, Pratt SC, Pavlic TP, Walker SI. 2020 Revealing the structure of information flows discriminates similar animal social behaviors. Elife 9, e55395. (doi:10.7554/eLife.55395)

14. Mizumoto N, Dobata S. 2019 Adaptive switch to sexually dimorphic movements by partner-seeking termites. Sci. Adv. 5, eaau6108. (doi:10.1126/sciadv.aau6108)

15. Matsuura K, Nishida T. 2001 Comparison of colony foundation success between sexual pairs and female asexual units in the termite Reticulitermes speratus (Isoptera: Rhinotermitidae). Popul. Ecol. 43, 119–124. (doi:10.1007/PL00012022)

16. Mizumoto N, Yashiro T, Matsuura K. 2016 Male same-sex pairing as an adaptive strategy for future reproduction in termites. Anim. Behav. 119, 179–187. (doi:10.1016/j.anbehav.2016.07.007)

17. Matsuura K, Kuno E, Nishida T. 2002 Homosexual tandem running as selfish herd in Reticulitermes speratus: novel antipredatory behavior in termites. J. Theor. Biol. 214, 63–70. (doi:10.1006/jtbi.2001.2447)

18. Li G, Zou X, Lei C, Huang Q. 2013 Antipredator behavior produced by heterosexual and homosexual tandem running in the termite Reticulitermes chinensis (Isoptera: Rhinotermitidae). Sociobiology 60, 198–203. (doi:Doi: 10.13102/sociobiology.v60i2.198-203)

19. Matsuura K, Fujimoto M, Goka K. 2004 Sexual and asexual colony foundation and the mechanism of facultative parthenogenesis in the termite Reticulitermes speratus (Isoptera, Rhinotermitidae). Insectes Soc. 51, 325–332. (doi:10.1007/s00040-004-0746-0)

20. Bartumeus F, Catalan J. 2009 Optimal search behavior and classic foraging theory. J. Phys. A Math. Theor. 42, 434002. (doi:10.1088/1751-8113/42/43/434002)

21. Leuthold RH. 1977 Postflight communication in two termite species, Trinervitermes bettonianus and Hodotermes mossambicus. In Proc. VIII Congr. IUSSI, pp. 62–64.

22. Bordereau C, Pasteels JM. 2011 Pheromones and chemical ecology of dispersal and foraging in termites. In Biology of Termites: A Modern Synthesis (eds DE Bignell, Y Roisin, N Lo), pp. 279–320. Dordrecht: Springer Netherlands. (doi:10.1007/978-90-481-3977-4_11)

23. Ampion M, Quennedey A. 1981 The abdominal epidermal glands of termites and their phylogenetic significance. Syst. Assoc. Spec. Vol. 19, 249–261.

24. Bailey NW, French N. 2012 Same-sex sexual behaviour and mistaken identity in male field crickets, Teleogryllus oceanicus. Anim. Behav. 84, 1031–1038. (doi:10.1016/j.anbehav.2012.08.001)

25. Gross MR. 1996 Alternative reproductive strategies and tactics: diversity within sexes. Trends Ecol. Evol. 2, 92–98.

26. Mason RT, Crews D. 1985 Female mimicry in garter snakes. Nature 316, 59–60. (doi:10.1038/316059a0)

27. Wendelken PW, Barth RH. 1985 On the significance of pseudofemale behavior in the neotropical cockroach genera Blaberus, Archimandrita and Byrsotria. Psyche (New York) 92, 493–503. (doi:10.1155/1985/97012)

28. Luo C, Wei C. 2015 Intraspecific sexual mimicry for finding females in a cicada: Males produce ‘female sounds’ to gain reproductive benefit. Anim. Behav. 102, 69–76. (doi:10.1016/j.anbehav.2015.01.013)

29. Thornhill R. 1979 Adaptive female-mimicking behavior in a scorpionfly. Science (80-.). 205, 412–414. (doi:10.1126/science.205.4404.412)

30. Gosden TP, Svensson EI. 2009 Density-dependent male mating harassment, female resistance, and male mimicry. Am. Nat. 173, 709–721. (doi:10.1086/598491)

31. Kuriwada T. 2017 Male–male courtship behaviour, not relatedness, affects the intensity of contest competition in the field cricket. Anim. Behav. 126, 217–220. (doi:10.1016/j.anbehav.2017.02.009)

32. Lane SM, Haughan AE, Evans D, Tregenza T, House CM. 2016 Same-sex sexual behaviour as a dominance display. Anim. Behav. 114, 113–118. (doi:10.1016/j.anbehav.2016.01.005)

33. MacFarlane GR, Blomberg SP, Kaplan G, Rogers LJ. 2006 Same-sex sexual behavior in birds: expression is related to social mating system and state of development at hatching. Behav. Ecol. 18, 21–33. (doi:10.1093/beheco/arl065)

34. Burgevin L, Friberg U, Maklakov AA. 2013 Intersexual correlation for same-sex sexual behaviour in an insect. Anim. Behav. 85, 759–762. (doi:10.1016/j.anbehav.2013.01.017)

35. Mizumoto N, Dobata S. 2018 The optimal movement patterns for mating encounters with sexually asymmetric detection ranges. Sci. Rep. 8, 3356. (doi:10.1038/s41598-018-21437-3)

36. Reynolds AM. 2006 Optimal scale-free searching strategies for the location of moving targets: New insights on visually cued mate location behaviour in insects. Phys. Lett. A 360, 224–227. (doi:10.1016/j.physleta.2006.08.047)

37. Pfau D, Jordan CL, Breedlove SM. 2021 The de-scent of sexuality: Did loss of a pheromone signaling protein permit the evolution of same-sex sexual behavior in primates? Arch. Sex. Behav. 50, 2267–2276. (doi:10.1007/s10508-018-1377-2)

38. Grosjean Y, Grillet M, Augustin H, Ferveur JF, Featherstone DE. 2008 A glial amino-acid transporter controls synapse strength and courtship in Drosophila. Nat. Neurosci. 11, 54–61. (doi:10.1038/nn2019)

39. Mitaka Y, Akino T. 2021 A Review of Termite Pheromones: Multifaceted, Context-Dependent, and Rational Chemical Communications. Front. Ecol. Evol. 8. (doi:10.3389/fevo.2020.595614)

40. Regaiolli B, Sandri C, Rose PE, Vallarin V, Spiezio C. 2018 Investigating parental care behaviour in same-sex pairing of zoo greater flamingo (Phoenicopterus roseus). PeerJ 2018. (doi:10.7717/peerj.5227)

41. Yamanaka O, Takeuchi R. 2018 UMATracker: An intuitive image-based tracking platform. J. Exp. Biol. 221, 1–24. (doi:10.1242/jeb.182469)

42. R Core Team. 2020 R: A language and environment for statistical computing.

43. Mizumoto N, Rizo A, Pratt SC, Chouvenc T. 2020 Termite males enhance mating encounters by changing speed according to density. J. Anim. Ecol. 89, 2542–2552. (doi:10.1111/1365-2656.13320)

44. Mizumoto N, Lee S Bin, Valentini G, Chouvenc T, Pratt SC. 2021 Coordination of movement via complementary interactions of leaders and followers in termite mating pairs. Proc. R. Soc. B Biol. Sci. 288, 20210998. (doi:10.1098/rspb.2021.0998)

45. Couzin ID, Krause J, James R, Ruxton GD, Franks NR. 2002 Collective memory and spatial sorting in animal groups. J. Theor. Biol. 218, 1–11. (doi:10.1006/jtbi.2002.3065)

46. Warton DI, Hui FKC. 2011 The arcsine is asinine: The analysis of proportions in ecology. Ecology 92, 3–10. (doi:10.1890/10-0340.1)

47. Bartumeus F, Levin SA. 2008 Fractal reorientation clocks: Linking animal behavior to statistical patterns of search. Proc. Natl. Acad. Sci. U. S. A. 105, 19072–19077. (doi:10.1073/pnas.0801926105)

48. Schielzeth H et al. 2020 Robustness of linear mixed-effects models to violations of distributional assumptions. Methods Ecol. Evol., 2041–210X.13434. (doi:10.1111/2041-210x.13434)

49. Mizumoto N, Bourguignon T. 2021 The evolution of body size in termites. Proc. R. Soc. B Biol. Sci. 288, 20211458. (doi:10.1098/rspb.2021.1458)

50. Sillam-Dussès D, Sémon E, Lacey MJ, Robert A, Lenz M, Bordereau C. 2007 Trail-following pheromones in basal termites, with special reference to Mastotermes darwiniensis. J. Chem. Ecol. 33, 1960–1977. (doi:10.1007/s10886-007-9363-5)

51. Revell LJ. 2012 phytools: An R package for phylogenetic comparative biology (and other things). Methods Ecol. Evol. 3, 217–223. (doi:10.1111/j.2041-210X.2011.00169.x)

